# Developmental senescence orchestrates hyaloid vessel regression in the postnatal eye

**DOI:** 10.64898/2026.05.12.724389

**Authors:** Alexa Silva Sosa, Agnieszka Dejda, Gaelle Mawambo, Gael Cagnone, Kawtar Zouine, Roberto Diaz, Vera Guber, Frédérick A. Mallette, Jean-Sebastien Joyal, Przemyslaw Sapieha, Malika Oubaha

**Affiliations:** Department of Biological Sciences, Faculty of Sciences, Université du Québec à Montréal (UQAM); Centre de Recherches sur les Maladies Orphelines - Fondation Courtois (CERMO-FC); Department of Ophthalmology, Maisonneuve-Rosemont Hospital Research Centre, University of Montreal, Montreal, Quebec, H1T 2M4, Canada; Department of Biochemistry and Molecular Medicine, University of Montreal, Maisonneuve-Rosemont Hospital Research Centre, Montreal, Quebec, H1T 2M4, Canada; Departments of Pediatrics, and Pharmacology, Centre Hospitalier Universitaire Ste-Justine Research Center, Montréal, Quebec, H3T 1C5 Canada; Department of Medicine, University of Montreal, Maisonneuve-Rosemont Hospital Research Centre, Montreal, Quebec, H1T 2M4, Canada

**Keywords:** hyaloids, developmental senescence, single cell RNA sequencing, vascular pruning, persistence vitreous

## Abstract

The mammalian eye develops in concert with coordinated growth and remodeling of three vascular networks: the hyaloid vasculature, the choroid and retinal plexus. While retinal and choroidal systems support visual function in the mature eye, the hyaloid network plays a vital yet temporary role supporting the developing lens and inner retina. Regression of the hyaloid network is essential for optical clarity, yet the mechanisms guiding the process remain incompletely understood.

Using single-cell RNA sequencing, we show that postnatal mouse hyaloid cells are broadly senescent. Hyaloid vascular smooth muscle, endothelial and immune cells display cell-cycle arrest marked by Cdkn1a with the expression of SASP factors. Genetic ablation of *Cdkn1a* impedes normal hyaloid regression, demonstrating that developmental senescence is essential for vascular remodeling and functions alongside apoptosis and macrophage-mediated clearance. These findings identify an unrecognized senescence-driven mechanism orchestrating hyaloid involution during ocular development, broadening the understanding of vascular remodeling in the eye.

## INTRODUCTION

The eye depends on sustained metabolic support throughout development. Before formation of the retinal vascular plexus, the transient hyaloid vasculature provides oxygen and nutrients to the primary vitreous, embryonic lens and retina. This vascular network subsequently regresses to permit formation of a clear optical path required for vision[1]. The hyaloid system originates from the hyaloid artery (HA) and comprises three interconnected components: the vasa hyaloidea propria (VHP), branching over the posterior lens surface; the tunica vasculosa lentis (TVL), covering the posterior lens and connecting with the choroidal vasculature; and the pupillary membrane (PM), located on the anterior lens surface[1, 2].

In humans, defective hyaloid vessel regression leads to Persistent Fetal Vasculature (PFV), a congenital anomaly characterized by the retention of embryonic vascular structures within the vitreous. Typically, PFV is associated with fibrovascular proliferation, lens opacities and in severe cases, tractional retinal detachment, often leading to irreversible vision impairment[1-5]. Studies on PFV and hyaloid regression identified that involution of the fetal hyaloid vasculature is a tightly regulated process triggering several overlapping signaling networks[6-9]. During late human embryogenesis, the hyaloid vasculature undergoes programmed regression, enabling the maturation of the vitreous and establishment of the retinal vasculature[10].

In mice, regression of hyaloid vasculature begins at birth and continues until approximately postnatal day (P) 21[11]. This process is primarily driven by apoptosis of endothelial cells, coupled with macrophage-mediated clearance to facilitate vessels regression[5]. Wnt7b, a Wnt ligand, produced by macrophages in proximity of hyaloid vessels, activates Wnt signaling by binding frizzled 4 (FZD4)/LRP5 receptors in adjacent endothelial cells to initiate apoptosis and subsequently vessel regression[5]. A role for neuronal control of hyaloid regression has been proposed with retinal neuron-derived vascular endothelial growth factor (VEGF) hyaloid vessels maintenance through VEGF receptor 2 (VEGFR2) signaling, with its depletion disrupting the balance and leading to persistent vasculature[7]. Similarly, melanopsin, a photoreceptive protein, has been implicated in modulating hyaloid remodeling, suggesting a link between light perception and vascular regression[12].

Cellular senescence is a multifaceted process characterized by a stable growth arrest and activation of tumor suppressor pathways such as Cdkn2a/Rb and p53/Cdkn1a(p21)[13-16]. Growth arrest associated to senescent cells can be triggered in pathological conditions by various stimulus including oncogenic signaling, DNA-damage, nutrient deprivation, oxidative stress, and chemotherapeutic drugs[17-21]. Subsequently, hallmarks of cellular senescence include elevated lysosomal activity, chromatin remodeling, metabolic reprogramming, autophagy, and secretion of bioactive molecules, a process called the senescence associated secretory phenotype (SASP)[22-28]. The SASP is a mixture of pro-angiogenic and pro-inflammatory cytokines, chemokines, growth factors, and proteases that reinforces cell cycle arrest, modifies the extracellular matrix and promotes the recruitment of immune cells to eliminate senescent cells[29-32]. The paracrine effect of SASP also plays beneficial roles in promoting wound healing and neuronal survival[26, 33]. Programs of cellular senescence are also widespread throughout developing embryonic tissues from mice, chicks and human[34-37]. Relevantly, endothelial cell cycle arrest has been shown to contribute to hyaloid vessel involution by restricting proliferative capacity[4]. While a framework is starting to be built where cellular senescence, proliferation and cell death collectively orchestrate embryonic development, there remain gaps in our knowledge[35, 37].

In the present study, we identify cellular senescence as a feature of murine hyaloid vasculature remodelling and involution, and show that inhibition of Cdkn1a, a central mediator of cell-cycle arrest and developmental senescence, affects this process.

## RESULTS

### Senescence-Associated β-Galactosidase activity is detected in the embryonic mouse eye

To examine the spatiotemporal distribution of cellular senescence during embryogenesis, we performed whole-mount staining for senescence-associated β-galactosidase (SA-β-Gal) in C57BL/6 mouse embryos at embryonic day 14.5 (E14.5) and E18.5 (**Fig. 1a, b**). Consistent with previous work by Serrano and Keyes [35, 37], SA-β-Gal activity was detected at E14.5 in multiple regions of the embryos, including the limb, tail tip, and otic vesicle (**Fig. 1a**). By E18.5, staining had largely disappeared from these extraocular regions, whereas strong SA-β-Gal activity persisted in the eye (**Fig. 1b**), indicating that the developing ocular compartment remains a prominent site of senescence-associated activity during late gestation.

**Figure 1.**
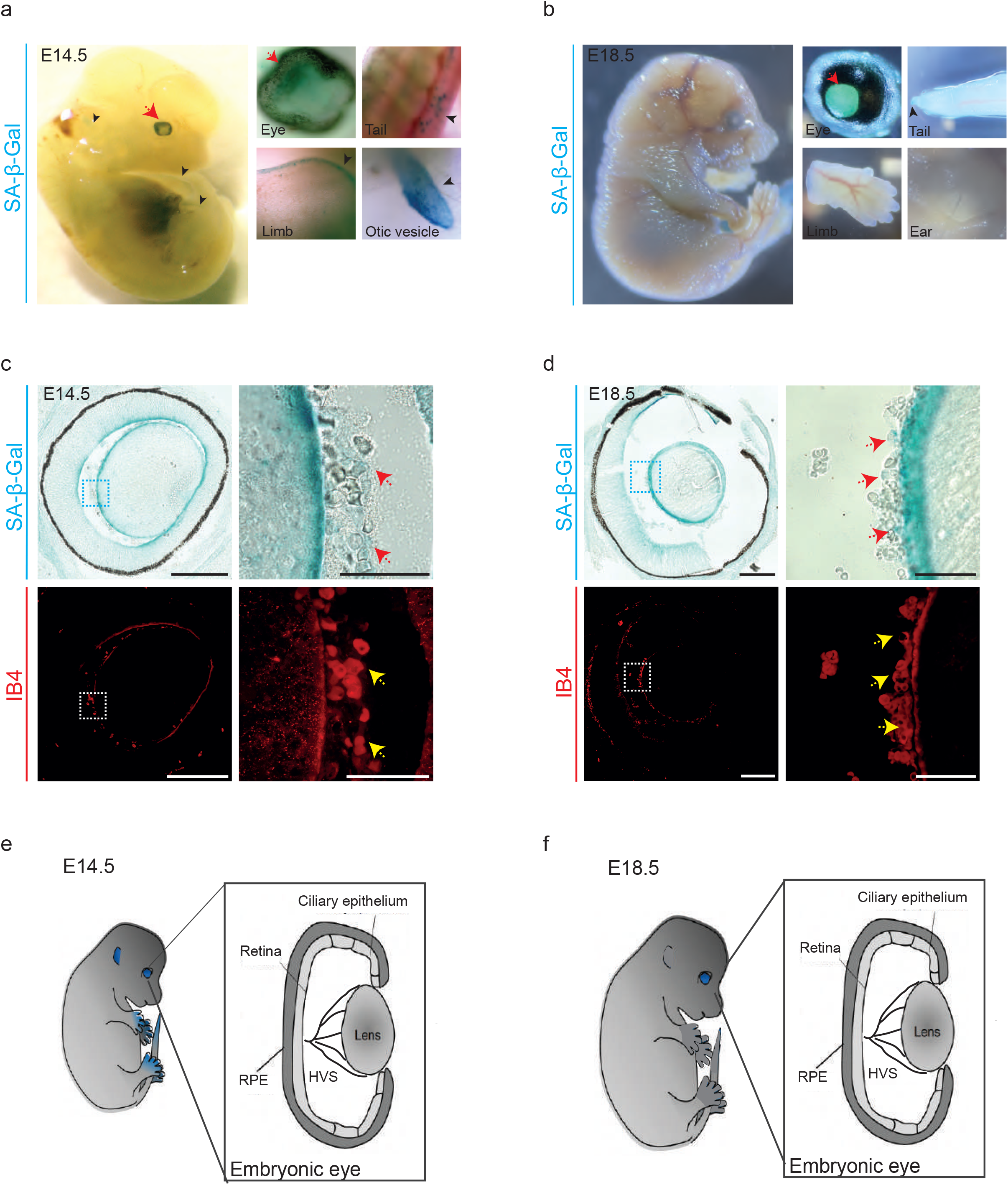
Senescence-Associated β-Galactosidase (SA-β-Gal) staining in mouse embryonic development. Whole C57BL/6 mouse embryos were stained for Senescence-Associated β-Galactosidase (SA-β-Gal). Representative images of stained positive regions are shown. **a**-Wholemount embryo showing strong SA-β-Gal staining in the eye, tail, limb, and otic vesicle at embryonic days (E) 14.5 and **b-** in the eye and the tip of the tail at E18.5. **Red** arrows show the embryonic eye and **black** arrowheads point to other senescent embryonic tissues. **c**-SA-β-Gal and Isolectin B4 (IB4) staining on cryosections mouse eyes at E14.5 and **d-** E18.5. SA-β-Gal staining revealed positive structures at the surface of the lens and in the vitreous, between the retina and the lens and areas surrounding the lens. IB4 staining shows specifically the localization of the vasculature within the eye. Higher-magnification views of the outlined areas are shown. **Red** arrows show SA-β-Gal^+^ cells and **yellow** arrowheads indicate blood vessel localization. Scale bars, 200 µm and 50 µm [for higher magnification]. **e-** Representative scheme of the embryo showing features of senescence (**blue**) identified at E14.5 and **f-** E18.5. Higher-magnification of transverse sections of the embryonic eye (E14.5-E18.5) are shown. HVS: Hyaloid Vascular System, RPE: Retinal Pigment Epithelium.

To obtain cellular resolution of SA-β-Gal staining during ocular development, we stained sagittal cryosections of embryonic mouse eyes at E14.5 and E18.5 (**Fig 1c, d**). At both stages, SA-β-Gal-positive cells were detected along the lens surface and within discrete domains of the vitreous, notably in the space between the lens and the retina (**Fig. 1c, d**; top panels). Co-staining with isolectin B4 (IB4) revealed that these SA-β-Gal-positive cells were positioned within the hyaloid vascular network (**Fig. 1c, d**; bottom panels), supporting an association between senescence-associated activity and the developing hyaloid vasculature. The anatomical context of these observations is summarized schematically in **Fig. 1e, f**, which highlights the persistence of SA-β-Gal activity in the eye from mid-to late gestation and its spatial relationship with the hyaloid vascular system. Together, these data identify the embryonic mouse eye as a persistent site of senescence-associated β-galactosidase activity and indicate that this signal is associated with cells of the hyaloid vasculature.

### Developmental senescence marks the regressing hyaloid vasculature, but not the developing retinal vessels

Despite their close anatomical proximity, the hyaloid and retinal vascular beds follow distinct developmental trajectories and remain spatially segregated during remodeling. While retinal vasculature continues to expand and mature, the hyaloid vessels undergo progressive regression (**Fig. 2a**). To determine whether cellular senescence persists postnatally and the compartment-specific distribution of SA-β-Gal-positive cells, we analyzed flat-mounts of hyaloid and retinal tissues from the same postnatal (P) eye at different developmental stages P0, P4 and P8. SA-β-Gal staining was detected exclusively in the regressing hyaloid vasculature and was absent from the developing retinal vessels (**Fig. 2b, c**). Furthermore, we examined cryosections of murine eyes collected at P0, P4 and P8. In these sections, SA-β-Gal activity co-localized with IB4-positive hyaloid vessels within the vitreous and around the lens capsule (**Fig. S1a, b**), confirming that senescence-associated activity remains confined to the regressing hyaloid compartment during early postnatal development. Together, these findings identify the hyaloid vasculature as a major site of developmental senescence in the postnatal eye and further highlight the distinct remodelling programmes that distinguish the regressing hyaloid network from the expanding, non-senescent retinal vasculature.

**Figure 2.**
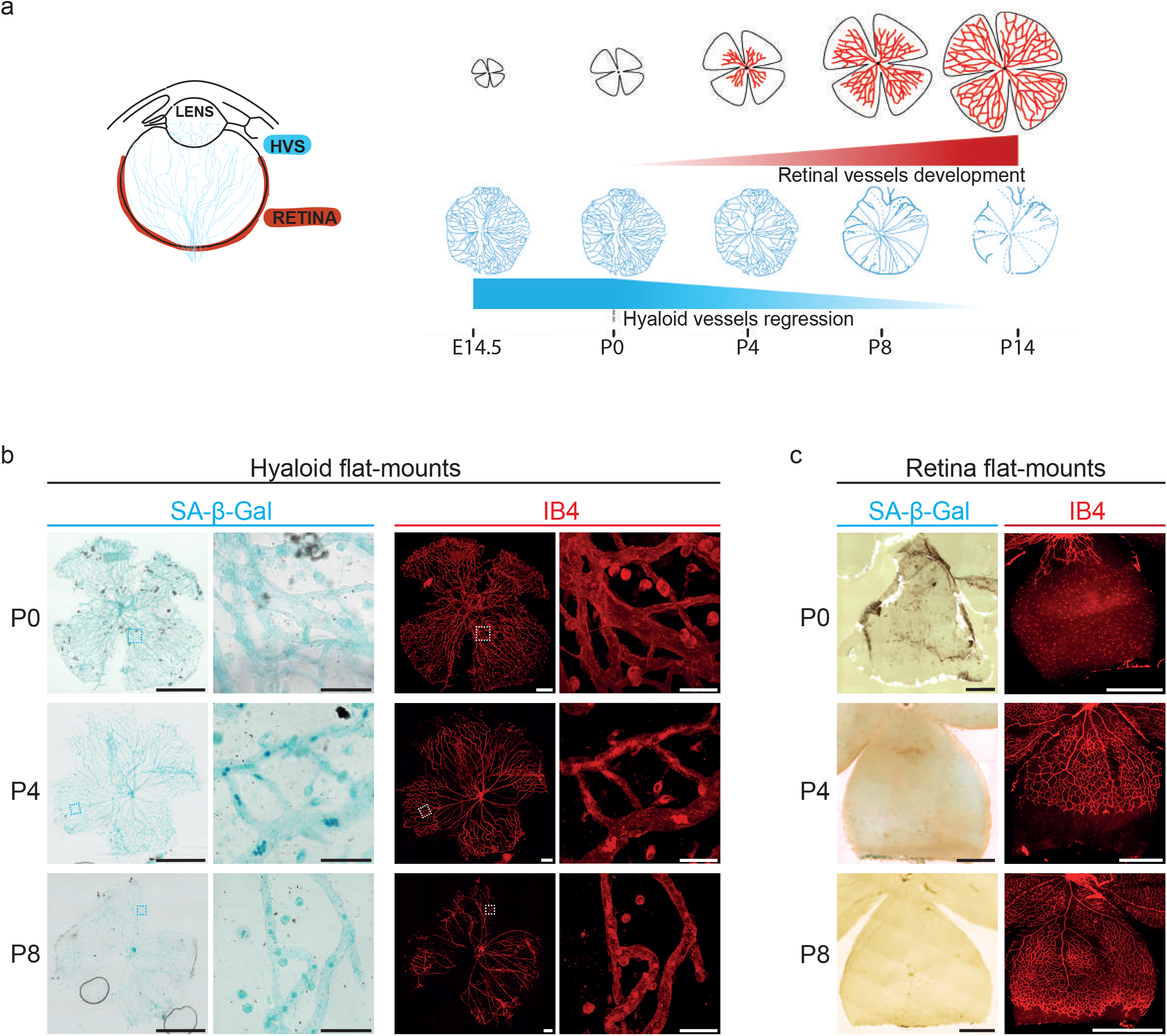
Restricted cellular senescence to hyaloid vessels during postnatal eye mouse development. **a-** Schematic illustration of mouse hyaloid vessels regression and retinal vascular growth from E14.5 to postnatal day (P)8. **b**- Confocal micrographs of representative SA-β-Gal and IB4 staining of hyaloid vessels and **c-** retina flat-mounts from the same mouse eye at P0, P4 and P8. Higher-magnification images of boxed hyaloid vessels areas are shown. Scale bars, 200 µm and 50 µm [for higher magnification].

### Transient fetal hyaloid vessels express molecular markers of cellular senescence

Selective SA-β-Gal staining in postnatal hyaloid vessels, with absence of signal in the developing retinal vasculature suggests distinct developmental programs **(Fig. 2)**. To further examine this difference at the molecular level, we compared published bulk RNA-seq data from P5 hyaloid vessels [38] to pseudo-bulk single-cell RNAseq at P5 retina[39]. This analysis revealed that hyaloids were enriched for pathways associated with cellular stress and senescence, including cytokine production, cytokine signaling, and senescence-related gene signatures. Hyaloids were also enriched for pathways involved in vascular remodeling and morphogenesis, including angiogenesis, endothelial cell migration, and epithelial-to-mesenchymal transition (**Fig. 3a**).

**Figure 3.**
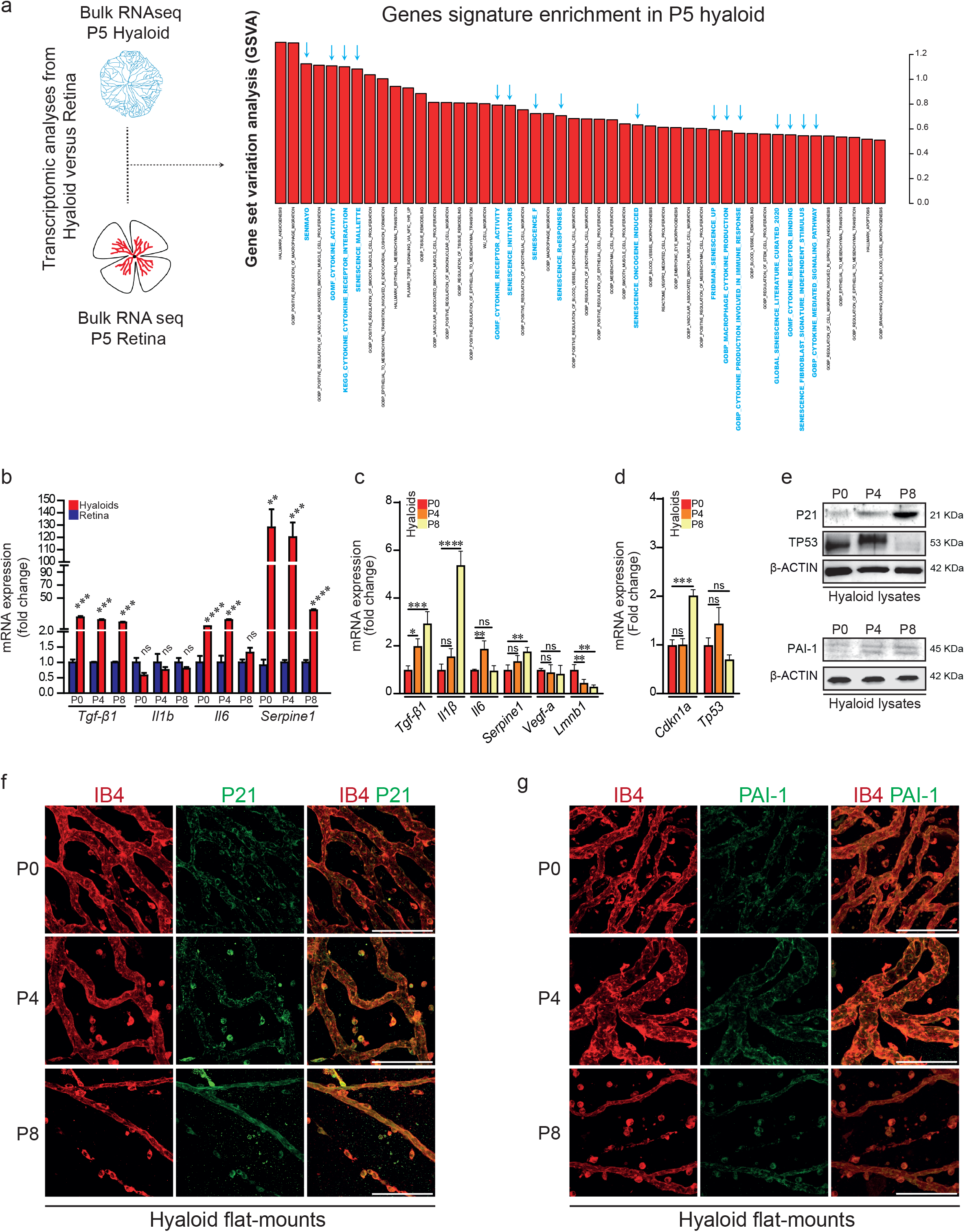
Enrichment of cellular senescence signature in hyaloid vessels. **a**-Transcriptome analyses from P5 hyaloid bulk-RNA sequencing data from R.A. Lang group compared to P5 retina pseudo bulk single-cell RNA sequencing data from S. Blackshaw group. Enrichment analysis was performed on selected mouse gene sets by Gene Set Variation Analysis (GSVA) and coefficient comparison was performed using limma with the difference in GSVA enrichment score (contrast fit coefficient) displayed as bar plot. **Blue** arrows point senescence-related pathways. **b-** Quantitative reverse transcription polymerase chain reaction (qRT-PCR) of SASP-related genes (*Tgf-β1, Il1β, Il6, Serpine1*) comparing between hyaloids and retina at P0, P4 and P8. *β-Actin* was used as reference gene. Data are presented as fold change compared to retina at each developmental stage (n=3). Bar graphs are means ± SEM. Represented *P* values are **<0.01, ***<0.001 and \*\*\*\**<*0.0001 from two-tailed parametric unpaired *t* -test. **c**- Bar charts of qRT-PCR from SASP-related genes (*Tgf-β1, Il1β, Il6, Serpine1, Vegf-a*) and **d-**cell cycle arrest markers (*Cdkn1a, Tp53*) in hyaloids at P0, P4 and P8. *Lmnb1* is used as a negative control. *β-Actin* was used as a reference gene. Data are presented as fold change compared to P0 (*n* ≥3-6). Bar graphs are represented as means ± SEM. Represented *P* values are *<0.05, **<0.01, ***<0.001 and ****<0.0001 from ordinary one-way ANOVA test followed by Dunnett’s multiple comparisons for *Tgf-β1, Il1β, Il6, Serpine1, Vegf-a* and *Cdkn1a* and Kruskal-Wallis test followed by Dunn’s multiple comparisons for *Tp53*. **e**- Immunoblots of hyaloid cell lysates from P0, P4 and P8 for senescence markers (P21, TP53, PAI-1). β-ACTIN was used as a loading control (n = 3). **f**- Representative confocal immunofluorescence staining of P21 (green) and **g**- PAI-1 (green) of flat-mounted hyaloid vessels at P0, P4 and P8. Blood vessels were stained with IB4 (red). Scale bars, 100 µm. Non-significant (ns).

To define this program more directly, we examined the expression of key senescence mediators and SASP factors in dissected hyaloid vessels and retinal tissue from postnatal mouse eyes. Compared with retina, hyaloids showed sustained upregulation of *Tgf-β1* and *Serpine1* at P0, P4, and P8, whereas *Il6* was significantly elevated at P0 and P4; in contrast, *Il1β* expression did not differ significantly between the two tissues (**Fig. 3b**). Within hyaloid vessels, mRNA transcripts of representative SASP factors, including *Tgf-β1, Il1β* and, *Serpine-1* were significantly upregulated at P8, and *Il6* was increased at P4 during hyaloid regression (**Fig. 3c**). In contrast, *Vegf-a* mRNA levels remained unchanged while *Lmnb1*, whose decrease is indicative of cellular senescence [40], declined significantly at P4 and P8 (**Fig. 3c**).

We next assessed the gene expression of key cell cycle arrest regulators: *Cdkn1a* and *Tp53*, in hyaloid vessels. C*dkn1a* exhibited a significant increase at P8, at both the mRNA and protein levels, while *Tp53* mRNA remained unchanged compared to TP53 protein that decreases at P8 compared to P0 (**Fig. 3d, e**). In addition, immunoblot analysis of postnatal hyaloid vessels confirmed expression of senescence marker PAI-1, a component of the SASP (**Fig. 3e**).

To map senescence within the regressing hyaloid, we assessed the spatial localization of p21, a canonical developmental senescence marker, together with PAI-1. Spatial expression of p21 and PAI-1 across defined postnatal stages revealed that both markers localize within IB4-positive blood vessel structures (**Fig. 3f, g**). P21 and PAI-1 were localized along vessel walls (likely endothelial cells) and in perivascular/extravascular compartments (likely mononuclear phagocytes). Collectively, these results suggest that pathways of cellular senescence are activated during programmed involution of the hyaloid vasculature.

### Single-cell RNA sequencing reveals distinct senescence signatures in hyaloid vascular populations

To elucidate the cellular composition and transcriptional programs underlying the programmed involution of postnatal hyaloid vessels, we performed single-cell RNA sequencing (scRNA-seq) on mouse hyaloid blood vessels isolated at P4 to capture transcriptomes of individual cells of hyaloid vasculature and resolve dynamic gene expression changes associated with vascular remodeling (**Fig. S2a**). This time point was selected as it coincides with a transitional window during hyaloid vessel regression marked by endothelial cell clearance[5], active vascular remodeling and early activation of stress- and senescence-related pathways (**Fig. 3a**).

Unbiased clustering using uniform manifold approximation and projection (UMAP) identified nine distinct cell populations within the P4 hyaloid vasculature dataset (**Fig. 4a**). Cell type annotation was guided by the top differentially expressed genes per cluster, ranked by mean expression and log2 fold change (log2FC) (**Fig. S2b**). Among the clusters we identified immune cells, endothelial cells, and vascular smooth muscle cells (VSMCs)/pericytes, canonical constituents of vasculature during development as shown by dot plot of gene markers (**Fig. 4a; S2b**). Six additional clusters were also detected, including lens precursors, epithelial cells, neural precursors, erythroblasts, corneal basal epithelial cells, and retinal ganglion cells (RGCs) as the hyaloid vessels are embedded within the neural retina (**Fig. 4a; S2b)**.

**Figure 4.**
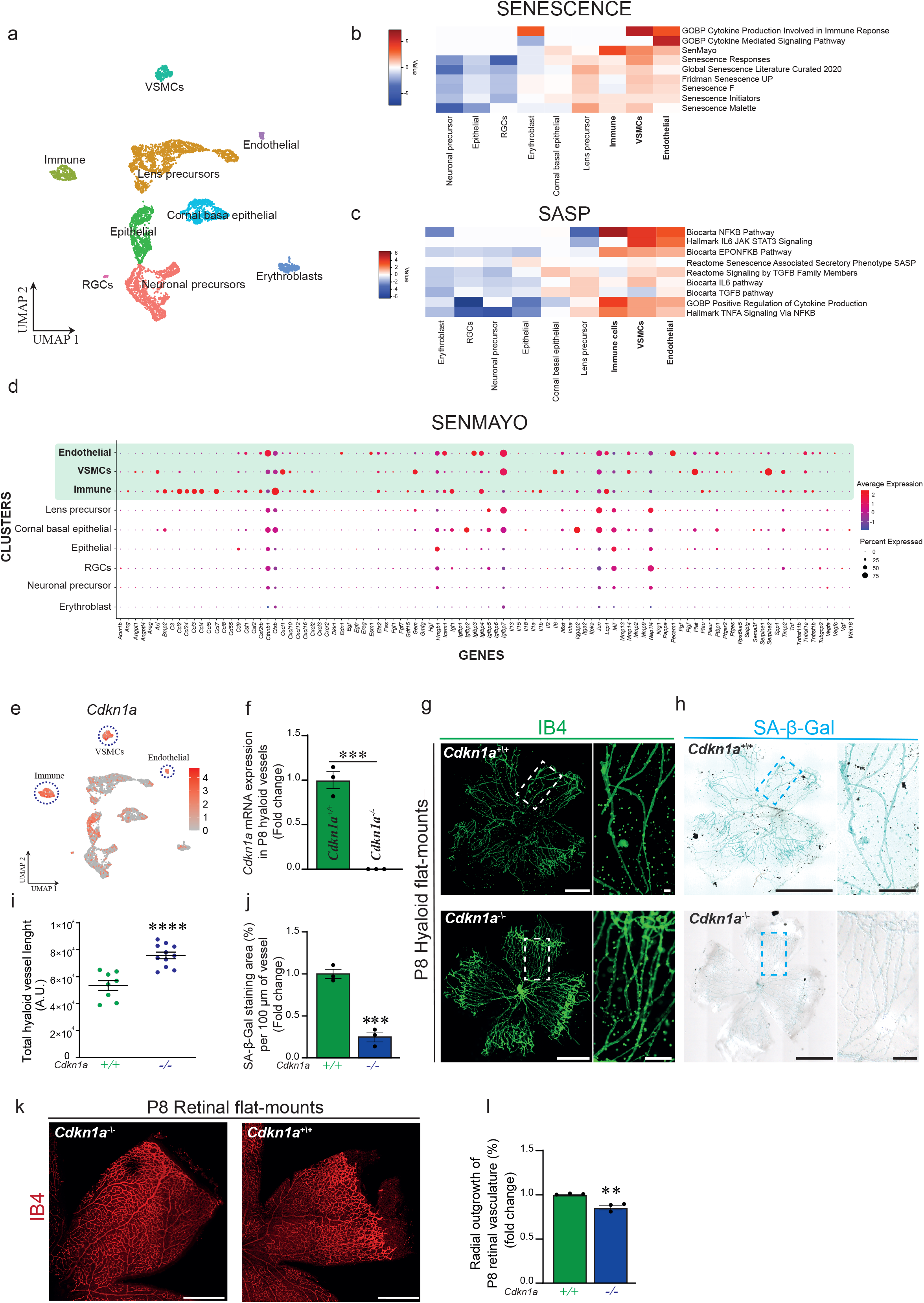
Single-cell analysis identifies cell-type-specific senescence programs and Cdkn1a-dependent regulation of hyaloid vessel regression. **a-** Uniform manifold approximation and projection (UMAP) of single-cell RNA sequencing from P4 hyaloid vessels representing the 9 hyaloid cell types identified by graph-based clustering of normalized RNA count. **b-** Heat map displaying log2 FC of VISION calculated enrichment scores across the 9 hyaloid cell populations for selected senescence- and **c**- SASP-related gene sets, show enrichment in immune, VSMCs and endothelial cell clusters. **d**- Dot plot illustrating the expression levels of SenMayo-related genes across the 9 hyaloid cell clusters. **e-** UMAP feature plot representation of *Cdkn1a* expression across the 9 hyaloid cell populations. **f**- Relative mRNA levels of *Cdkn1a* in P8 hyaloid vessels *Cdkn1a*^*+/+*^ versus *Cdkn1a*^*-/-*^ measured by qRT-PCR (n = 3). *β-Actin* was used as reference gene. **g-** Representative IB4 and **h-** SA-β-Gal staining of flat-mounted P8 hyaloid vessels *Cdkn1a*^*+/+*^ (control) and *Cdkn1a*^*-/-*^ (lacking Cdkn1a). Higher magnifications of vessels are shown. **i**- Quantification of total vessel length of P8 flat-mounted hyaloids *Cdkn1a*^*+/+*^ control versus *Cdkn1a*^*-/-*^. Each dot represents a flat-mounted hyaloid sample (n = 8-11). **j-** Quantification of SA-β-Gal staining per 100 µm of vessels (in percentage) of P8 *Cdkn1a*^*+/+*^ versus *Cdkn1a*^*-/-*^ flat-mounted hyaloid. **k-** Representative IB4 staining of P8 retinal flat-mount *Cdkn1a*^*+/+*^ and *Cdkn1a*^*-/-*^ and **l**- quantification of radial outgrowth of retinal vascular plexus (n = 3), showing delayed expansion of the superficial vascular plexus in *Cdkn1a*^*-/-*^ mice. Data are presented as fold change normalized to *Cdkn1a*^*+/+*^ control (**f, j, l**) or as Arbitrary Unit (A.U.) (**i**). Data are represented as means ± SEM (**f, j, l**) or as individual values (**i**). Represented *P* values are \**\**< 0.01, *\*\*\**< 0.001 and *\*\*\*\**< 0.0001 from two-tailed parametric unpaired *t* -test. Scale bars, 500 µm (**g, h**) and 100 µm (**k**) [for higher magnification in **g, h**].

These findings are consistent with prior single-cell RNA sequencing of mouse hyaloid tissues at P3 and P6, which similarly identified various non-vascular cell types, including astrocytes, melanocytes, and fibroblasts, within the vitreous compartment[41]. Given our focus on hyaloid regression, we restricted subsequent analyses to the three vascular relevant populations: endothelial cells, immune cells, and VSMCs/pericytes (**Fig. S2c**).

To examine the heterogeneity of the senescence program within hyaloid blood vessels, we assessed the enrichment of senescence-associated gene sets across hyaloid cell populations at P4. A scaled comparison of gene and pathway expressions between hyaloid populations at P4 shows that immune, VSMCs and endothelial cells, each exhibited distinct senescence and secretory signatures (**Fig. 4b-d**). Notably, and consistent with the SA-β-Gal signal at the lens periphery (**Fig. S1a**), senescence-related gene sets were enriched in lens precursor population. Together, these transcriptional data support the induction of canonical senescence markers and SASP factors in hyaloid vessels detected by immunoblotting and qRT-PCR (**Fig. 3b-d**).

P4 single-cell RNA-sequencing analysis further identified activation of cell-death programmes alongside developmental senescence (**Fig. S2d**). In line with this, cleaved caspase-3 was broadly detected across the regressing hyaloid from P0 to P8 by immunofluorescence (**Fig. S2e**).

Developmental senescence is tightly regulated by the cyclin-dependent kinase inhibitor Cdkn1a, underscoring the intersection of cell cycle regulation and developmental patterning[35, 37]. Given that Cdkn1a is abundantly expressed across multiple cell populations within the hyaloid vasculature in the eye (**Fig. 4e)**, we investigated rates of hyaloid regression in mice lacking Cdkn1a (*Cdkn1a*^*-/-*^) (**Fig. 4f**), which are described as developmentally normal[42, 43]. Hyaloid vessels (IB4 staining) were considerably more elaborate, persisted longer and showed a significant reduction in SA-β-Gal intensity at P8 in Cdkn1a-deficient mice (*Cdkn1a*^*-/-*^) compared to the wild-type control (*Cdkn1a*^*+/+*^) (**Fig. 4g-j)**. To determine whether hyaloid vessel persistence in *Cdkn1a*-deficient mice affects retinal vascular development, we performed IB4 staining on flat-mounted retinas. Loss of Cdkn1a led to delayed expansion of the superficial vascular plexus, indicative of impaired angiogenic progression (**Fig. 4k, l**).

These findings suggest coordinated remodeling processes at this stage, and it will be of particular interest to investigate how these programs evolve at earlier and later postnatal stages. Together, our data indicates that hyaloid cell populations display distinct and temporally regulated senescence programs, suggesting coordinated remodeling processes during developmental regression. Moreover, they suggest a contribution of Cdkn1a and by extension developmental senescence in the remodeling of hyaloid vasculature.

## DISCUSSION

This study identifies a previously unrecognized role for programmed cellular senescence in mouse ocular vascular morphogenesis, establishing senescence as a component of vascular regression in the transient hyaloid vasculature. In adult tissues, senescence is typically governed by the coordinated action of multiple cell-cycle regulators, including p16^Ink4a^, p19^Arf^, p53, and p21 [44-47]. Our data, together with previous studies, support a role for Cdkn1a in developmental senescence, while not excluding contributions from other regulators [34, 35, 37]. Consistent with this, Cdkn1a is expressed in the postnatal hyaloid vasculature, and its deletion reduces senescence and is associated with persistence of the hyaloid vessels, supporting a role for p21-dependent developmental senescence in hyaloid vessel remodelling. Together, these findings extend developmental senescence to the transient ocular vasculature, beyond previously described contexts such as the limb, mesonephros and placenta[34, 35, 37].

Furthermore, senescence-associated features were evident from embryonic stages and persisted throughout postnatal hyaloid development, preceding clear morphological signs of vessel involution. This temporal pattern is consistent with the possibility that senescence contributes to the developmental regression programme, rather than arising as a secondary consequence of tissue involution. More broadly, these findings align with the view that developmental senescence is a tightly regulated and context-dependent process, engaged with precise spatiotemporal control during the remodelling and elimination of transient structures such as the hyaloid vasculature, placenta and mesonephros[34, 35].

The persistence of senescence markers beyond embryogenesis and throughout postnatal stages points to a distinctive regulatory programme in the eye, unlike other embryonic tissues in which senescence has been identified. Notably, the senescence-associated secretory phenotype of hyaloid vessels differs substantially from that described in other embryonic senescent tissues[35, 37]. Hyaloid senescence was associated with strong expression of Il-6, Il-1β, Tgf-β1, and Pai-1, consistent with local inflammatory priming, extracellular matrix remodelling and immune recruitment. As proper hyaloid regression is essential for normal ocular development, our findings may have relevance for congenital eye disease and vitreoretinopathies linked to persistent fetal vasculature. It will therefore be important to examine the dynamics of cellular senescence in mouse models of hyaloid persistence, including *Ndp*^*-/-*^ or *Frizzled-4*^*-/-*^ (*Fzd-4*^*-/-*^)[48, 49].

More broadly, previous studies in p63, PASG and BRCA1 mutant models have shown aberrant embryonic senescence[50-52], raising the possibility that disruption of developmental senescence may contribute more widely to genetic disorders affecting early eye development.

Our data reveals that hyaloid senescence programs are both cell-type specific and developmentally dynamic, suggesting that physiological senescence can play a beneficial role in normal vascular regression.

In conclusion, we identify a Cdkn1a-associated senescence programme in the developing hyaloid vasculature, defined by cell-type-specific senescence states and a distinct SASP signature. Our work positions senescence as a developmentally relevant and instructive component of hyaloid regression, acting alongside apoptosis to coordinate vascular remodelling. Endothelial cells may either undergo apoptosis or enter senescence, while macrophages likely mediate clearance of apoptotic and senescent material to ensure orderly vessel elimination. These findings broaden the framework of eye vascular remodelling and suggest that targeting senescence or clearance pathways could offer new avenues to correct persistent hyaloid vessels in PFV.

### Limitations of the Study

While our findings reveal a previously unrecognized role for Cdkn1a-driven senescence in hyaloid vessel regression, several limitations warrant consideration. First, although SA-β-Gal and p21 staining provided spatial and temporal resolution of senescent cells, we were unable to definitively assign these markers to specific cell types *in situ*, limiting our understanding of cell-type-specific senescence dynamics. Single-cell RNA sequencing offered transcriptional insights, but functional validation of a role for cellular senescence in each hyaloid cell population remains to be established throughout postnatal stages of eye development. Additionally, our genetic ablation model focused solely on Cdkn1a; thus, potential contributions of other senescence regulators, such as p53, p16 or p19, cannot be excluded. These limitations highlight the need for additional studies with inducible and lineage-specific senescence reporters, to dissect the spatiotemporal complexity of developmental senescence in these transient blood vessels of the eye.

## MATERIAL AND METHODS

### Mice

Mice procedures were performed according to the Association for Research in Vision and Ophthalmology Statement for the Use of Animals in Ophthalmic and Vision Research and were approved by the Animal Care Committee of the University of Montreal in agreement with the guidelines established by the Canadian Council on Animal Care. C57BL/6 wild-type and Cdkn1a (p21) (*Cdkn1a*^*tm1Led*^/J, no. 016565) knock-out (*Cdkn1a*^*-/-*^)[43] mice lines were obtained from The Jackson Laboratory. Both female and male littermates were used, but sex information was not collected.

### Western blot

For assessment of hyaloid protein levels, eyes were enucleated from mice at different developmental stages, postnatal day (P) 0, P4 and P8. RIPA buffer with anti-protease and anti-phosphatase (BioRad) was freshly prepared to manually with a piston to homogenize the tissues. Protein concentration was determined by using a bicinchoninic acid (BCA) protein assay kit (23227; ThermoFischer Scientific) and 50 µg of cell lysates were analyzed for each time-point by standard SDS-PAGE technique. Primary antibodies used for immunoblotting are listed in **Table S1**. Secondary antibodies dependent on the produced species were used (#1706515, #1706516; BioRAD) at a dilution 1:3000.

### Immunofluorescence

To localize the ocular vasculature, mice eyes were enucleated at embryonic days (E)13.5 and E18.5 and postnatal stages P0, P4 and P8, and fixed in 4 % paraformaldehyde (PFA) (043368-9M; ThermoFischer Scientific) (1 hour, room temperature (RT)) followed by incubation in 30 % sucrose (16 hours, 4 °C). The whole eyes were embedded in OCT (Optimal Cutting Temperature) compound (23-730-571; Fisher Healthcare) at -20 °C and performed 10-µm sections. Slices were incubated in PBS (pH=7.4) (10010023; Gibco) containing 1 % Triton X-100 (TB0198; Bio Basic), 3 % Bovine Serum Albumin (BSA), 3 % Fetal Bovine Serum (FBS) (090-450; Wisent) and 0.01 % sodium deoxycholate (1 hour, RT) and then stained with Griffonia Simplicifolia Lectin (GSL) I-B4 DyLight649 (Isolectin B4(IB4)) (DL-1208-.5; Vector Laboratories) (dilution 1:200) (1 hour, RT). Coverslips were mounted using Fluoromount (00495802; Sigma-Aldrich).

For specific visualization of panretinal and hyaloid vasculature, mouse eyeballs at P0, P4 and P8 were fixed in 4 % PFA (1 hour, RT) before stored in PBS (4°C). After removal of the sclera, the retinal cup (with hyaloid vessels and crystal body) was incubated (1 hour, RT) in PBS containing 1 % Triton X-100, 10 % FBS, and 10 mM glycine, and then stained with IB4 (1 hour, RT). The stained retinal cup was fixed in 4 % PFA (20 minutes, RT), soaked in 2 % low melting temperature (LMP) agar in PBS (37°C, 18 hours) followed by incubation at 4°C for 3 hours. The crystal body was removed, and the retinal layer and LMP-agar embedded hyaloid vessels were flat-mounted using Fluoromount.

For specific protein localization, hyaloid vessels were stained with specific primary antibodies (**Table S2**) and IB4. Secondary antibody Alexa Fluor 488 donkey anti-rabbit IgG (H+L) (A-21206; Invitrogen) (dilution 1:500) was used for detection.

We carried out immunofluorescence experiments using a Nikon A1/A1R confocal microscope. Images were assembled using Photoshop CS4 (Adobe Systems). The hyaloid vascular density was determined using ImageJ[53] with adapted methodology previously described for the retina vasculature[54].

### Gene set enrichment analysis (Retina P5 versus Hyaloid P5)

RNAseq gene expression counts of murine P5 hyaloid tissue was obtained from Nayal et al. Development 2018 (GEO acc. Number GSE113294) using the deposited FPKM count table (GSM3103009_Hyaloid_P4.5_FPKM_0.3.xlsx). Transcript FPKM counts were aggregated (sum) by gene, generating a P5 RNAseq hyaloid dataset. PseudoBulk gene expression counts of murine P5 retina tissue was obtained from Clark et al. Neuron 2029 (GEO acc. Number GSE118614) using the deposited single-cell RNAseq matrix (GSE118614_10x_aggregate.mtx.gz by 10X Genomics). The single-cell RNAseq matrix was loaded into Seurat V5[55] and only P5 data were subset, then RNA counts were averaged using the AverageExpression function. To reflect P5 RNAseq hyaloid, averaged counts from the P5 retinal single-cell RNAseq data were multiplied by 100, generating a pseudo-bulk P5 retina dataset. Both P5 hyaloid and retinal dataset were merged, duplicated or non-overlapping gene names were removed, and the resulting data frame was transformed into an ExpressionSet file using Biobase (http://www.nature.com/nmeth/journal/v12/n2/full/nmeth.3252.html). The expressionSet was then processed with GSVA[56] using pathway gene sets from the msigdb.v2025.1.Hs.symbols.gmt database as well as custom gene sets. GSVA results were analysed by limma package[57], making contrast between P5 hyaloid and P5 retina data followed by lmFit and contrasts.fit to obtain coefficients that were then plotted using barplot for selected gene sets.

### Single-cell transcriptomic data analysis (Droplet Sequencing or Drop-seq)

Droplet sequencing (Drop-seq) procedure was done on hyaloid vessel cells suspension isolated from mice at P4. Briefly, two biological replicates single-cell suspensions were prepared from 18 total eyes from 9 mice, through successive steps of digestion (using papain solution; LK003150, Worthington,), trituration, and filtration. Droplet generation and cDNA libraries were performed as described in the Drop-seq procedure[58], and sequencing was done on Illumina NextSeq 500. We obtained a total of 2824 and 2836 sequenced cells respectively.

Briefly, RNA counts for the two replicates were merged into one single Digital Gene Expression (DGE) matrix and processed using Seurat package[59]. Cells expressing less than 100 genes and more than 10% of mitochondrial genes were filtered out. Single-cell transcriptomes were normalized using SCtransfrom (vars.to.regress = c (“nFeature_RNA”, “percent.mt”). PCA analysis on the most variable genes in the DGE matrix identified 20 significant PCs, which served as input for Uniform Manifold Approximation and Projection (UMAP). We used a density clustering approach to identify putative cell types on the embedded map and computed average gene expression for each identified cluster based on Euclidean distances. We then compared each cluster to identify marker genes differentially expressed across clusters, which with previous known cell type markers allowed cluster annotation. Seurat visualization tools included Dot Plot and UMAP Plot. Single-cell gene expression profiles from specific cell types identified by scRNA-seq were further analyzed using the Pathway activity score tool VISION[60] and differences between cell types were calculated using the FindMarker function from Seurat, with log2FC represented on heatmaps using heatmap.2 from gplots (Warnes G, Bolker B, Bonebakker L, Gentleman R, Huber W, Liaw A, Lumley T, Maechler M, Magnusson A, Moeller S, Schwartz M, Venables B, Galili T (2025). *gplots: Various R Programming Tools for Plotting Data*. R package version 3.3.0, https://talgalili.github.io/gplots/).

### Senescence Associated-β-Galactosidase (SA-β-Gal) assay

Senescence Associated-β-Galactosidase (SA-β-Gal) assay was carried out as previously described [26]. Mouse samples (whole eye sections and retinas and hyaloids flat-mounted) were incubated in 1 mM Magnesium chloride (MgCl_2_)/PBS (pH=5.0) (18 hours, 4°C) followed by a solution containing 1 mg/mL X-Gal (16495; Cayman Chemical), 6.5 mM Potassium ferricyanide, 5 mM Potassium ferrocyanide and 1 mM MgCl_2_/PBS (pH=5.0) (4-8 hours, 37°C). We then rinsed briefly with PBS (pH=7.4) (10010023; Gibco) and mounted with Fluoromount (00495802; Sigma-Aldrich) for visualization under a bright-field microscope (Nikon color DS-Fi1).

### Quantitative reverse transcription polymerase chain reaction (qRT-PCR)

Total RNA from hyaloid vessels and retina tissues at varying time points (P0, P4, P8) was isolated using PureZol (#7326890; BIO-RAD) according to the manufacturer’s protocol. For qRT-PCR, cDNA samples were prepared from 1 µg of total RNA following the manufacturer’s protocol (AD100-31404; Diamed). Quantitative PCR was performed with 2x Universal SYBR Green Fast qPCR Mix (RK21203; ABClonal) on the Bio-Rad CFX96 Real-Time System (Hercules, CA, USA) and software. Melt curve analysis was conducted to confirm the specificity of the PCR products. The relative quantity of the mRNA was calculated using the comparative CT method. The expression levels of target genes were normalized to the levels of *β-Actin* as reference gene. The specific mouse primers pairs used are listed in **Table S3**.

### Quantification of SA-β-Gal *in vivo*

SA-β-Gal staining in flat-mounted hyaloids was analyzed using ImageJ software described in [26].

### Statistical analyses

Statistical analyses were performed using GraphPad Prism 8.0.2 (GraphPad Software, San Diego, CA). Data are presented as mean ± SEM or as individual values, as indicated. Normal data distribution was assessed using the Shapiro–Wilk normality test with a significance level of 0.05. For comparisons between two groups, two-tailed parametric unpaired *t*-tests were used in a normal distributed data. For comparisons involving more than two groups, ordinary one-way ANOVA followed by Dunnett’s multiple-comparison test was used for normally distributed data. When data distribution did not follow normality, the non-parametric Kruskal-Wallis test followed by Dunn’s multiple-comparison correction was applied. It was considered statistically significant if the *P* values are *<0.05, **<0.01, ***<0.001 and ****<0.0001. N for each condition indicates the number of experimental replicates.

**Table S1.** Primary antibodies used for Western blot analysis.

| Target | Clone | Company | Catalogue no. | Dilution |
| --- | --- | --- | --- | --- |
| TP53 | FL-393 | Santa Cruz Biotechnology | sc-6243 | 1/500 |
| P21 | C-19 | Santa Cruz Biotechnology | sc-397 | 1/500 |
| **PAI-1** | H-135 | Santa Cruz Biotechnology | sc-8979 | 1/500 |
| **β-ACTIN** |  | MEDIMABS | MM-0164-P | 1/2000 |

**Table S2.** Primary antibodies used for Immunofluorescence.

| Target | Company | Catalogue no. | Dilution |
| --- | --- | --- | --- |
| **P21** | ABClonal | A1483 | 1/100 |
| **PAI-1** | ABClonal | A19096 | 1/100 |
| **CLEAVED**<br>**CASPASE-3** | Cell Signaling | 9664S | 1/300 |

**Table S3.**
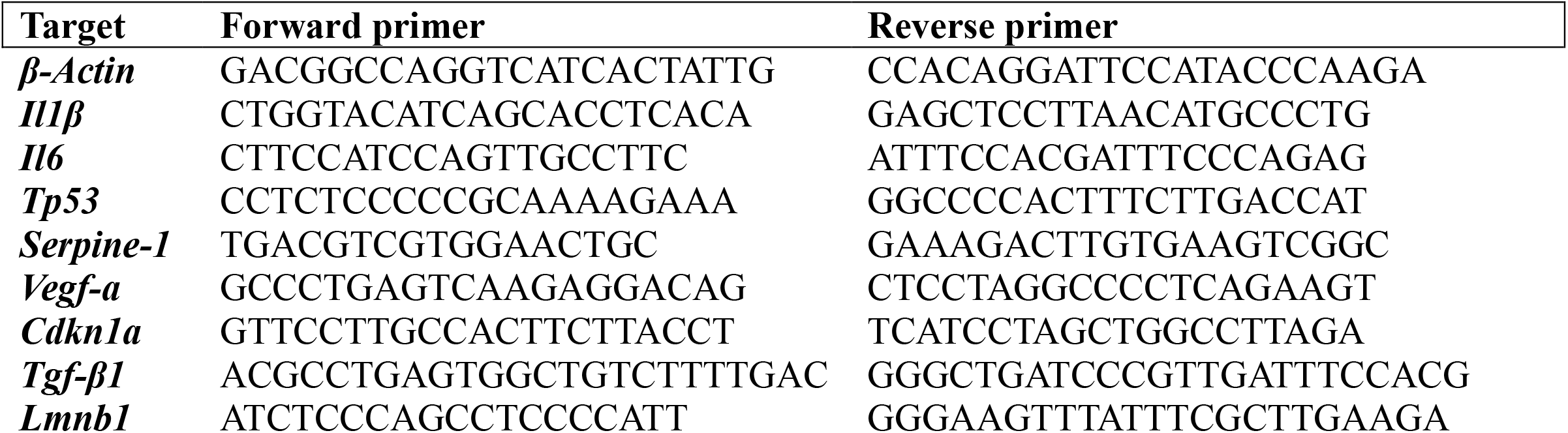
Table of primers used for qRT-PCR reagents of murine origin.

## Supplementary Figure legends

**Figure S1.**
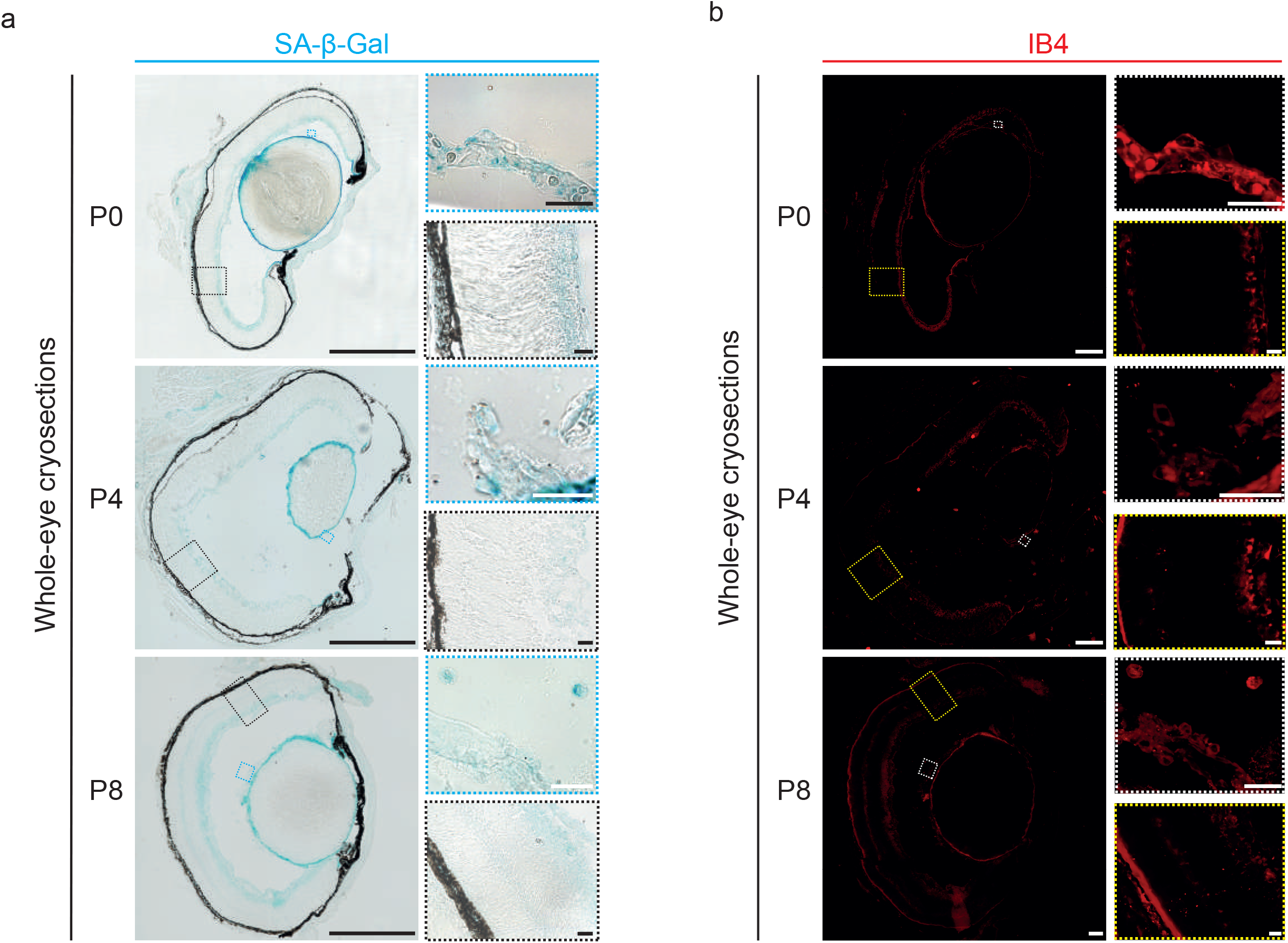
Whole eye SA-β-Gal staining reveals senescence in the postnatal mouse eye. **a-** Representative SA-β-Gal and **b**- IB4 staining of cryosections of mouse eyes at postnatal (P) stages P0, P4 and P8. Higher-magnification views of the outlined areas are shown. Scale bars, 200 µm and 25 µm [for higher magnifications].

**Figure S2.**
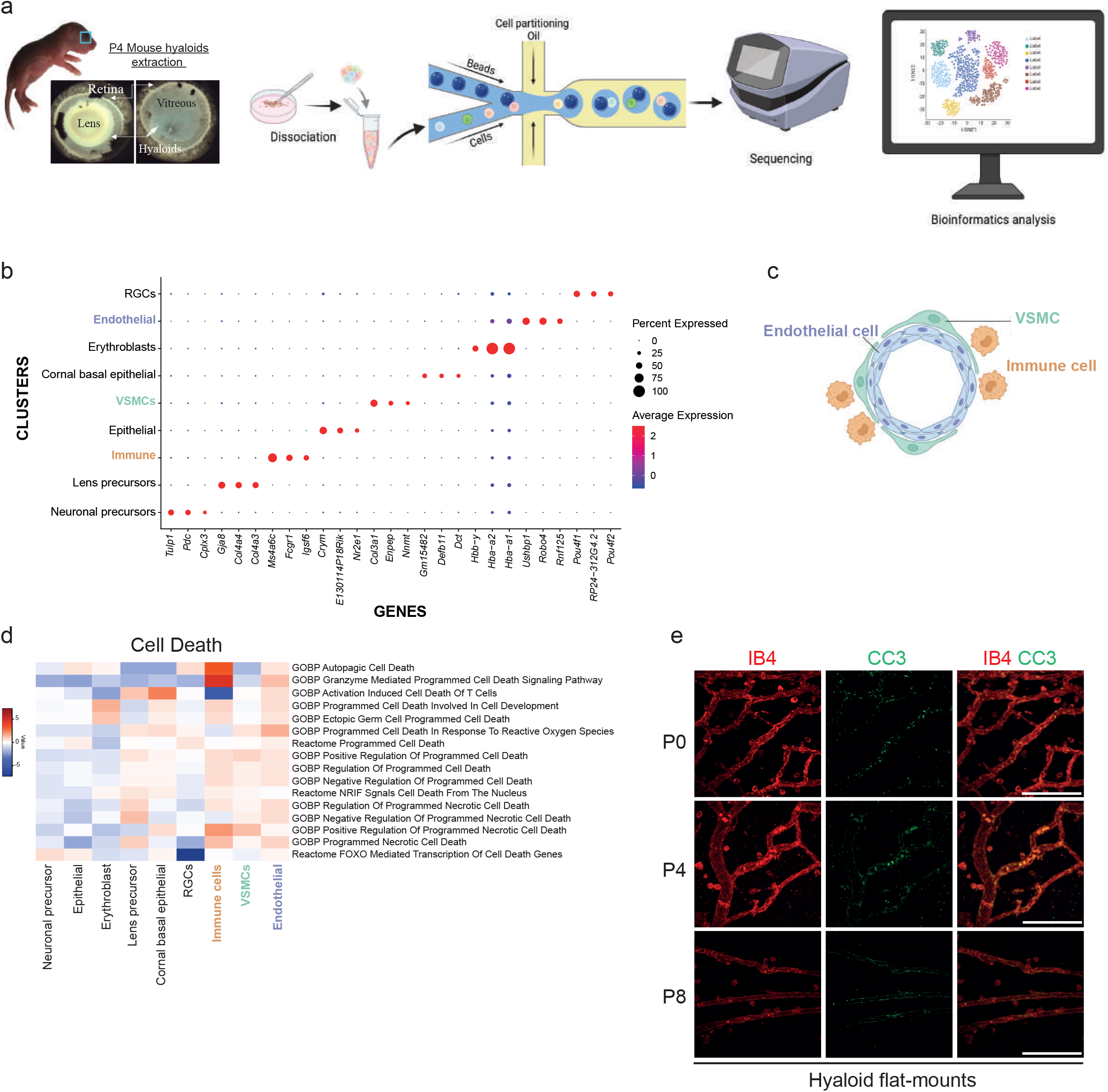
Cell death dynamics in hyaloid vessels during regression. **a**-Schematic illustration explaining the distinct phases of single-cell RNA sequencing analysis on dissected mice hyaloid vessels at P4. **b**- Dot plot of the top 3 marker genes for each of 9 hyaloid cell population identified. **c**- Graphical representation of the main component of the vascular unit of the hyaloid vessel. **d-** Heat map displaying log2 FC of VISION calculated enrichment scores across the 9 hyaloid cell populations for selected cell death-related gene sets. **e**- Representative confocal immunofluorescence staining of Cleaved-Caspase-3 (CC3) (green) of flat-mounted hyaloid vessels at P0, P4 and P8 (n=3). Blood vessels were stained with IB4 (red). Scale bars, 100 µm.

